# Synthetic cell-based tissues for bottom-up assembly of artificial lymphatic organs

**DOI:** 10.1101/2024.10.01.616088

**Authors:** Anna Burgstaller, Erick Angel Lopez Lopez, Oskar Staufer

## Abstract

Synthetic cells have emerged as novel biomimetic materials for studying fundamental cellular functions and enabling new therapeutic approaches. However, replicating the structure and function of complete tissues as self-organized 3D collectives has remained challenging. Here, we engineer lymph node-mimicking 3D lymphatic bottom-up tissues (lymphBUTs) with mechanical adaptability, metabolic activity, and hierarchical microstructural organization based on individual synthetic cells. We demonstrate that primary human immune cells spontaneously infiltrate and functionally integrate into these synthetic lymph nodes to form living tissue hybrids. By tuning the lymphBUT micro-organization and metabolic activity, we induce the *ex vivo* expansion of therapeutic CD8^+^ T cells with an IL-10^+^/IL-17^+^ regulatory phenotype. Our study highlights the functional integration of living and non-living matter, advancing synthetic cell engineering toward 3D tissue structures.

## Introduction

Tissues are hierarchical cellular architectures formed from numerous different cell types with varying functionality. The organization and compartmentalization of tissues underlays cellular physiology and supports regulation of intercellular signaling. Translating this degree of mesoscale cellular organization to lab-made biomaterials has remained a central ambition in the design of biomimetics of functional tissue-like structures [1]. In this realm, bottom-up synthetic biology assembles bionic cell-like materials with defined and controlled structural and functional properties. In most instances, microcompartments are assembled, reminiscent of living cells, in the form of individual and dispersed synthetic cells. Based on this recent progress in synthetic cell engineering, new opportunities arise to form whole synthetic cell-based tissues with advanced therapeutic functionalities.

While bottom-up assembly of dispersed synthetic cells with biomedical value remains a formidable engineering challenge, recent pre-clinical studies reported first successes of synthetic cells with therapeutic function, such as pro-angiogenic effects, stimulation of neuronal stem cell differentiation, targeted cancer cell killing and secretion of therapeutic proteins inside tumors [2–5]. Commonly applied synthetic cell models include unilamellar vesicles, coacervates, colloidosomes and microemulsions [6–8], each designed to mimic selected structural or functional features of living cells. In the context of synthetic tissue assembly, organized 3D constructs built from interconnect synthetic cell structures have been formed from water-in-oil emulsions [9]. Subsequently, bioprinting technologies were developed to form communicating 3D structures based aqueous droplet emulsions interconnected *via* lipid bilayers [10].

In a complementary approach, we previously development an oil-in-water-based dispersed synthetic cell model formed from an elastomer core surrounded by a lipid bilayer and functionalized with immune-stimulating ligands (at a density of 20 – 200 molecules/µm^2^) that can be interfaced with primary human immune cells [11]. This droplet supported lipid bilayer (dsLB) technology closely emulates the stiffness of natural T cell-activating dendritic cells (approx. 3 kPa) and presents membrane ligands with lateral mobility. Both stiffness and ligand mobility are critical factors for the formation of signaling competent membrane-membrane interfaces, as required for the formation of supramolecular adhesion architectures between cells [12]. Importantly, dsLBs can be cultured with human cells, activate primary human T cells *ex vivo* and expand therapeutically preferable phenotypes, making them an ideal synthetic cell models to explore for bottom-up synthetic tissue formation in an immunotherapeutic context.

Immunotherapy, specifically cell adoptive immunotherapy (CAI), is one of the most significant advancements of contemporary medicine. Typically, CAI is based on isolation of patient T cells, their expansion in an *ex vivo* environment and reinfusion into the patient after genetic modification [13]. The efficiency of CAI relies on controlled *ex vivo* expansion of therapeutic T cell phenotypes which is achieved by *ex vivo* T cell cultivation with artificial antigen presenting cells (aAPCs) in the form of solid dispersed polystyrene beads that mimic receptor activation of natural antigen presenting cells (APC). These are decorated with antibodies directed against CD3 to stimulate T cell receptor activation and CD28 acting as co-stimulatory signal. However, these dispersed, bead-based technologies do not fully emulate the *in vivo* 3D tissue context of lymph nodes. Lymph nodes are the physiological environment for T cell expansion, which has been shown to regulate the expansion behavior of T cells and contribute to their effector qualities.

For instance, recent studies have highlighted the relevance of the mechanical properties in the 3D microenvironment mimicking lymph nodes and therefore the controlled *ex vivo* T cell expansion [14–16]. This is highly relevant to simulate the naïve T cell recruitment, thus multiplying the lymph node mass two-to five folds and the increasing confinement and ECM stiffness as reaction to inflammation while maintaining nutrient supply [17]. Towards tissue engineering of a synthetic lymph node, Majedi *et. al* developed a 3D alginate-based porous hydrogel with tunable rigidity and equipped with anti CD3/CD28 antibodies. Using this model, they showed varying activation dynamics of T cells induced by different scaffold rigidities and suggested that higher confined scaffolds lead to increased T cell activation [11]. However, such approaches mostly recapitulate the lymph node extracellular matrix (ECM) and do not integrate living APCs or aAPCs. The combination of tissue mechanics and APC biochemistry is central for immune cell expansion. It therefore needs to be considered with the same degree of segmented, hierarchical organization found in lymph nodes to provide a relevant artificial model.

Towards engineering synthetic cell-based tissues with structured microanatomy of specialized biochemical and biomechanical functionalities, we here integrated dsLBs and vesicles to self-assemble into artificial lymphatic organs. These advanced tissue mimetics integrate biophysical properties and biochemical functionalities of natural APCs in a 3D tissue-like context. These lymphatic bottom-up tissues (lymphBUTs) can provide primary human T cells with a synthetic cellular environment that supports their expansion upon activation. Functional and structural heterogeneity naturally observed in lymphatic tissues are integrated to improve the efficiency of expansion and the quality of the differentiated T cells. LymphBUTs therefore present the first synthetic cell-based tissues, formed by self-assembly from cell-like bionic materials that sustain and develop the therapeutic functionality of human cells.

## Results

### LymphBUT formation and characterization

Towards efficiently forming synthetic cell-based 3D tissues with adjustable mechanical properties, a dsLB-based self-assembly process was implemented. The self-assembly mechanism relies on interconnecting individual dsLBs by the binding of biotin, integrated in the dsLB membrane *via* headgroup modified lipids, with tetrameric streptavidin (**Fig. 1 A**). Through continuous orbital shaking (to promote mixing and synthetic cell contact formation) in a physiological buffer, macroscale tissue-like structures can be assembled with sizes between 1.0 - 4.0 mm (**Fig. 1 B, Fig. 1 C**).

**Fig. 1:**
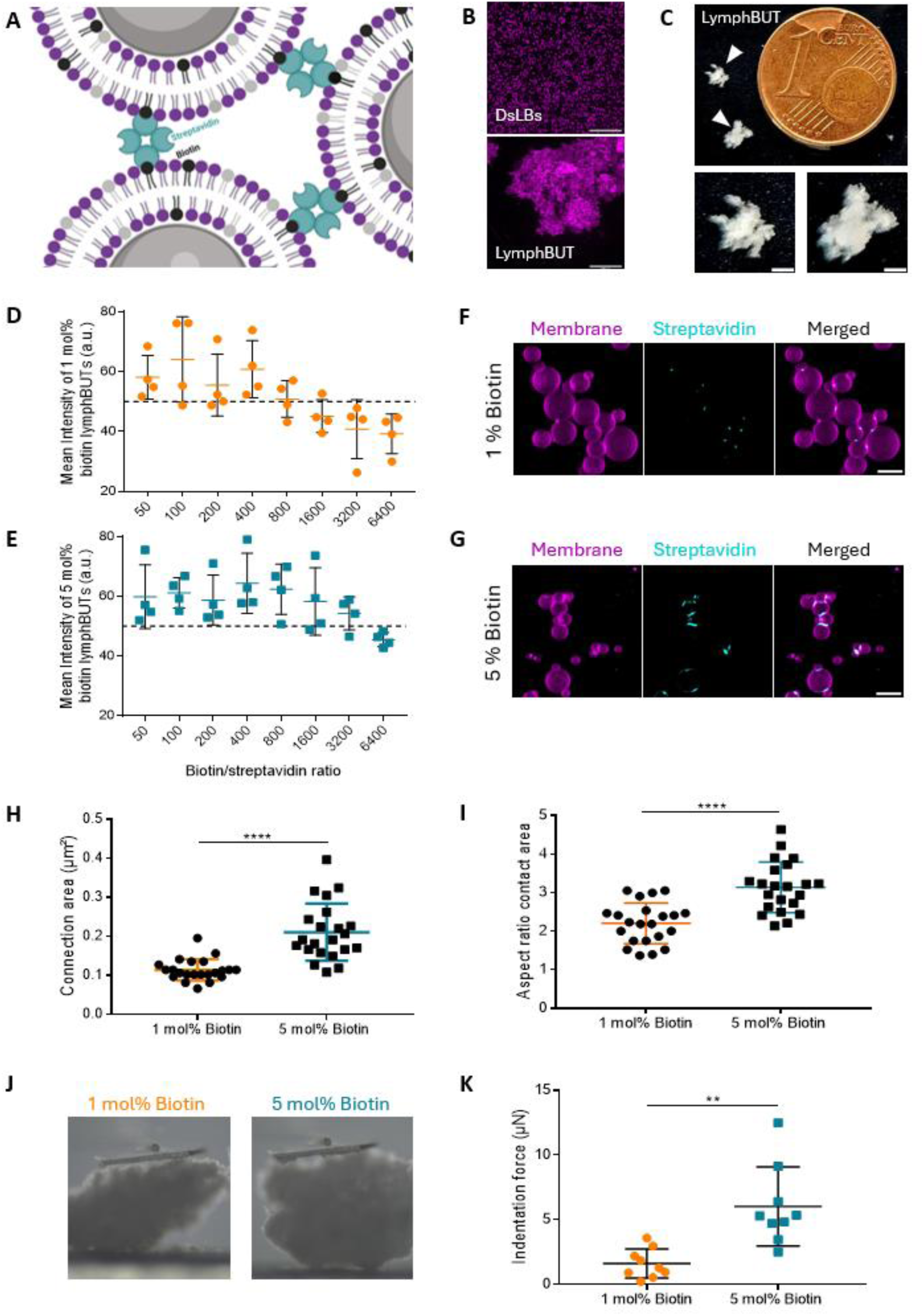
LymphBUT assembly and biomechanical characterization. **A)** Schematic illustration of the self-assembly process of dsLB-based synthetic cells into lymphBUT initiated by biotin-streptavidin interactions. **B)** Representative confocal microscopy maximum z-projections comparing disperse single dsLBs to a fully assembled lymphBUTs. Scale bars are 100 µm. **C)** Top view of two representative lymphBUTs (triangles) next to a 16.25 mm diameter 1 Eurocent for size comparison and lymphBUT magnifications. Scale bars are 1 mm. **D) and E)** Stereomicroscopy imaging based mean intensity quantification of lymphBUTs connected *via* 1 mol% (D) or 5 mol% (E) biotin in the dsLB membrane and streptavidin at varying biotin: streptavidin molecular ratios (50/100/200/400/800/1600/3200/6400:1). Dashed line indicates the threshold for fully formed lymphBUTs. The Mean intensity values are shown as mean +/- SD from n = 4 separate lymphBUT batches. **F) and G)** Representative confocal microscopy maximal z-projections of lymphBUTs built from dsLBs with 1 mol% (F) and 5 mol% (G) biotin in the membrane connected by AlexaFluor405-labeled streptavidin (1400:1), concentrating at the connection sides between the dsLBs. Scale bars are 10 µm. **H) and I)** Quantification of connection area (H) and connection aspect ratio (I) from images in F) and G). Results are shown as mean +/- SD from n = 22 manually segmented connection areas. P values were calculated using an unpaired tow-tailed student t test. **J)** Representative bright field images of 1 mol% and 5 mol% biotin lymphBUTs in the micro-indentation setup. **K)** Comparison of maximal indentation force required to compress 10 % of total height of lymphBUTs made from dsLBs with 1 mol% and 5 mol% biotin in the membrane. Results are shown as mean +/- SD from n = 9 lymphBUTs produced in two individual batches. P values were calculated using an unpaired tow-tailed student t test. ** p<0.01, **** p<0.0001

To investigate the impact of the streptavidin/biotin ratio on the lymphBUTs formation process and to potentially tune the mechanical properties of lymphBUTs, dsLBs with 1 mol% and 5 mol% biotin in the membrane were formed. These were added to solutions of varying streptavidin concentration between 50:1 and 6400:1 (biotin:streptavidin). Measuring the optical transmissibility to approximate lymphBUT density, we found that optimal ratios differ with varying biotin concentrations in the dsLB membrane (**Fig. 1 D, E**). DsLBs with 5 mol% biotin produced dens lymphBUTs at lower biotin/streptavidin ratios. This effect was observed over a broad range of streptavidin/biotin molecular ratio (50:1 to 3200:1). At ratios above 50:1 we observed streptavidin saturation of the dsLB membranes and therefore no or only fragile lymphBUTs formation (**Fig S. 1 A**). Ratios below 800:1 for 1 mol% and 3200:1 for 5 mol% biotin containing dsLBs did not show any lymphBUT formation. Of note, lymphBUTs could “self-heal” after mechanical rupture by prolonged incubation in up to at least 5 formation-destruction cycles without supplementation of further streptavidin (**Fig S. 1 B**).

We next investigated the microstructural differences between 1 mol% and 5 mol% lymphBUTs and imaged the connection areas between individual dsLBs by laser scanning confocal microscopy (LSCM) using AlexaFluor405-labeled streptavidin. In both cases, we found streptavidin enrichment at the contact sites between individual dsLBs, however, with varying structure (**Fig. 1 F, G**). In 1 mol% biotin lymphBUTs, spot-like connection clusters were observed, whereas 5 mol% biotin lymphBUTs showed significantly larger and more elongated planar connection areas (**Fig. 1 H, I**). This local enrichment also highlights the relevance of a laterally mobile membrane to sustain and support formation of the supramolecular connection sites and the relevance of cell deformability to allow for planar interaction sites. Towards assessing whether the dsLB elasticity also contributes to the lymphBUT formation process and contact site maturation, we test synthetic tissue formation with lipid membrane coated silica beads of comparable size (3.94 µm) and identical membrane composition. We observed aggregation and contact formation between the beads in the center of the culture well, induced by the orbital shaking, but no 3D synthetic tissue formation (**Fig S. 1 C**). This indicates that a mobile membrane is not sufficient for stable contact formation, but a deformable synthetic cell is required to form robust intercellular adhesion interfaces as also observed in natural tissues [18]. Therefore, synthetic tissue formation necessitates the deformability of synthetic cells allowing the formation of robust planar adhesive interfaces. From a macroscopic point of view, intercellular adhesion between cells in tissues impacts the overall tissue mechanics. To evaluate if different dsLB interaction architectures also translate to varying lymphBUT stiffnesses, we measured their compressibility by micro-indentation (**Fig.1 J**). We compared the maximum indentation force needed to indent 10 % of the total lymphBUTs height formed from 1 mol% and 5 mol% biotin dsLBs. The results demonstrate a significantly increased indentation force for 5 mol% biotin lymphBUTs compared to 1 mol% biotin lymphBUTs (**Fig. 1 K**), presumably based on the larger intercellular areas between dsLBs (**see Fig. 1 D, E**). Therefore, the lymphBUT tissue stiffness can be tuned by adjusting the inter-dsLB connection sites *via* varying biotin concentrations. This opens the door of synthetic cell-based tissues with adjustable stiffness and biochemical compositions properties.

### Heterotypic LymphBUT formation for primary human T cell activation

Towards activating and expanding T cells within lymphBUTs, we developed an approach for conjugating dsLBs with agonistic antibodies to induce T cell receptor activation (anti-CD3) and T cell co-stimulation (anti-CD28) [11]. DsLBs were produced with a maleimide modified lipid membrane (final concentration of 5 mol%) to couple antibodies to their surface (**Fig. 2 A**). LSCM imaging revealed a homogeneous distribution of AlexaFluor488-labeled antibodies on the dsLB membrane and across the population (**Fig. 2 B**). We calibrated the surface density of the antibodies on the dsLB membrane *via* beads of known molecules of equivalent soluble fluorochrome (MESF) (**Fig S. 2 A**). In further experiments, anti-CD3 and anti-CD28 antibodies were used at a final density of 420 molecules/µm^2^ (high) and 320 molecules/µm^2^ (low) in a 1:3 ratio. Importantly, when incubating CD8^+^ T cells with lymphBUTs, we observed spontaneous infiltration into the lymphBUT tissue. After 4 days of incubation, T cells formed confined activation clusters reminiscent of clonal expansion clusters in lymph nodes [19] (**Fig. 2 C**). We quantified the activated CD25^+^ T cells cultivated within lymphBUTs of different densities and antibody concentrations. Independently from the lymphBUT density and the antibody concentrations, the CD25 expression was between 32 % and 40 % of the total T cell population cultivated with lymphBUTs which is slightly lower as compared to the industry standard Dynabeads (**Fig. 2 D**). Interestingly, when quantifying the expression of the immune-checkpoint receptor PD-1 among the CD25^+^ T cells, we found that T cells expanded within lymphBUTs expressed a significantly reduced fraction of PD-1^+^ cells as compared to dispersed dsLBs and Dynabeads (**Fig. 2 E**). This indicates that the combination of biomechanical support provided by lymphBUTs efficiently mimics the confined activation process of T cells *in vivo* and opens the door to expand T cells with an improved phenotype which is less prone to immune suppression.

**Fig. 2.**
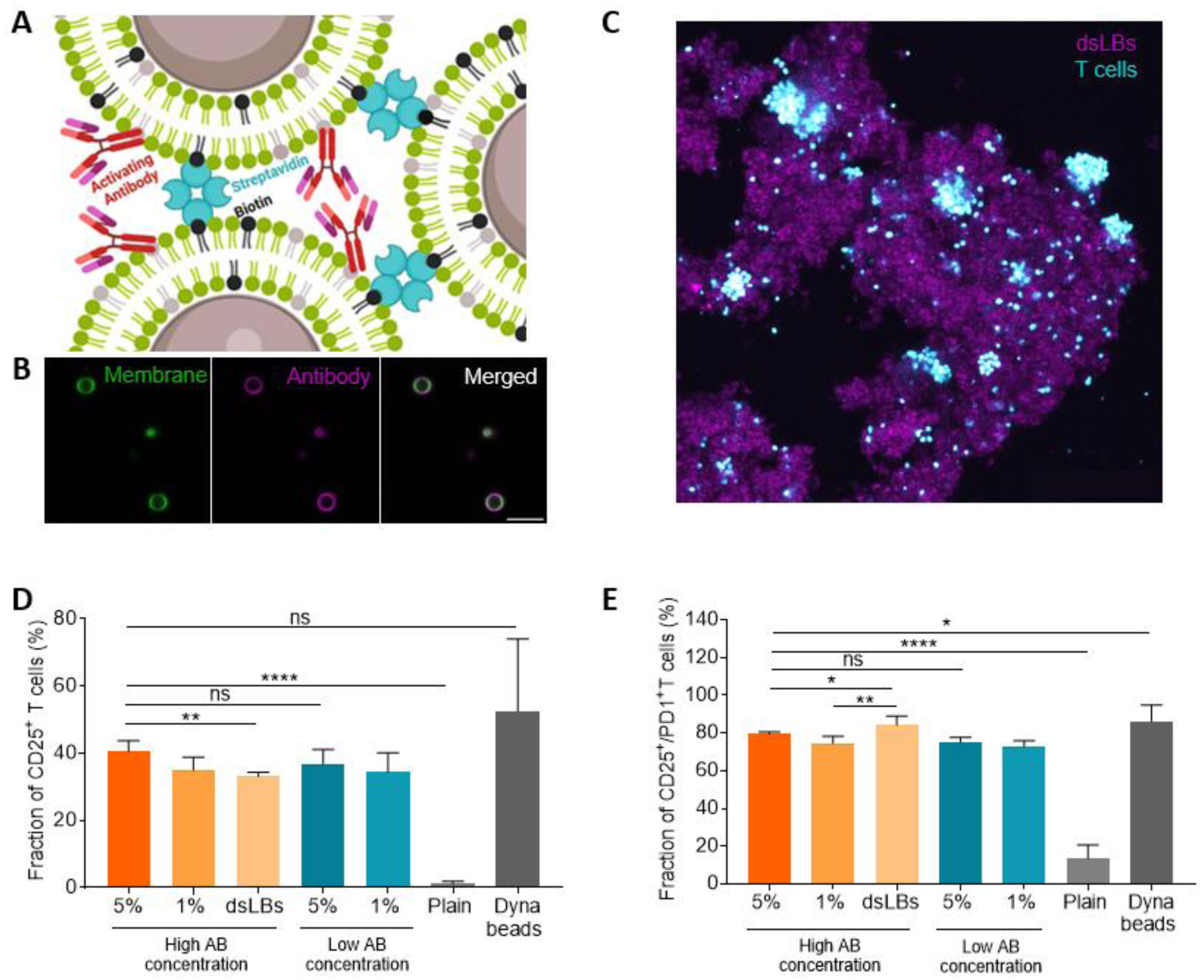
LymphBUT immune functionalization. **A)** Schematic illustration of immune-stimulating antibody decorated dsLBs interconnected by streptavidin and biotin. **B)** Representative confocal microscopy images of dsLBs functionalized with AlexFluor^TM^488-labeled goat anti-human IgG (magenta). Scale bar is 10 µm. **C)** Representative confocal microscopy maximal z-projections showing human T cells infiltrated into a 5 mol% biotin lymphBUT with a high anti-CD3 and anti-CD28 concentration. Scale bar 200 µm. **D) and E)** Quantification of the CD25 (D) and CD25/PD-1 (E) expressing T cell fraction expanded in different lymphBUT stiffnesses and antibody concentrations with dispersed dsLBs, Dynabeads or lymphBUTs which do not display antibodies (plain) measured by flow cytometry. Results are shown as mean +/- SD of two donors in n > 2 technical replicates. P values were calculated using two-tailed t test. ns = not significant p>0.05, * p<0.05, ** p<0.01, **** p<0.0001

Lymph nodes, like several lymphatic tissues, are organized in a hierarchical structure with spatially segregated tissue functions (e.g. T cell zones, germinal centers). This architecture is crucial to separate effector and regulatory cells and provides a mechanical environment for expansion. For instance, fibroblasts in the lymph node stroma provide mechanical support for lymph node swelling and controlled T cell expansion. Towards mimicking this multifunctional heterotypic cellular architecture with varying biochemical and biomechanical properties, we implemented a sequential formation process to build spatial heterogenous lymphBUTs (**Fig S. 2 B**) taking advantage of the lymphBUT self-healing properties (**Fig S. 1 B**). In a first step, lymphBUTs are formed from antibody-decorated, T cell-activating dsLBs and homogenized by resuspension into smaller lymphBUTs of approximately 50 - 100 µm size. In analogy to the dense dendritic cell network in T cell zones of lymph nodes, dsLBs with 5 mol% biotin were used providing the appropriate confinement. By subsequently adding non-functionalized dsLBs, a scaffold is formed surrounding the functionalized lymphBUTs. This loser network made of 1 mol% dsLBs is able to expand as a response to intra-lymphBUT T cell expansion. LSCM imaging confirmed formation of heterotypic lymphBUTs containing distinct stimulatory zones, surrounded by a non-stimulatory scaffold. We successfully formed heterotypic lymphBUTs with varying ratios of reaction zones to scaffold (75/25, 50/50, 25/75) showing segregated architectures as compared to homogeneous lymphBUTs formed from pre-mixed functionalized and non-functionalized dsLBs (**Fig. 3 A**). This demonstrates that the hierarchical architecture of natural tissues can be mirrored to a simplified degree.

**Fig. 3.**
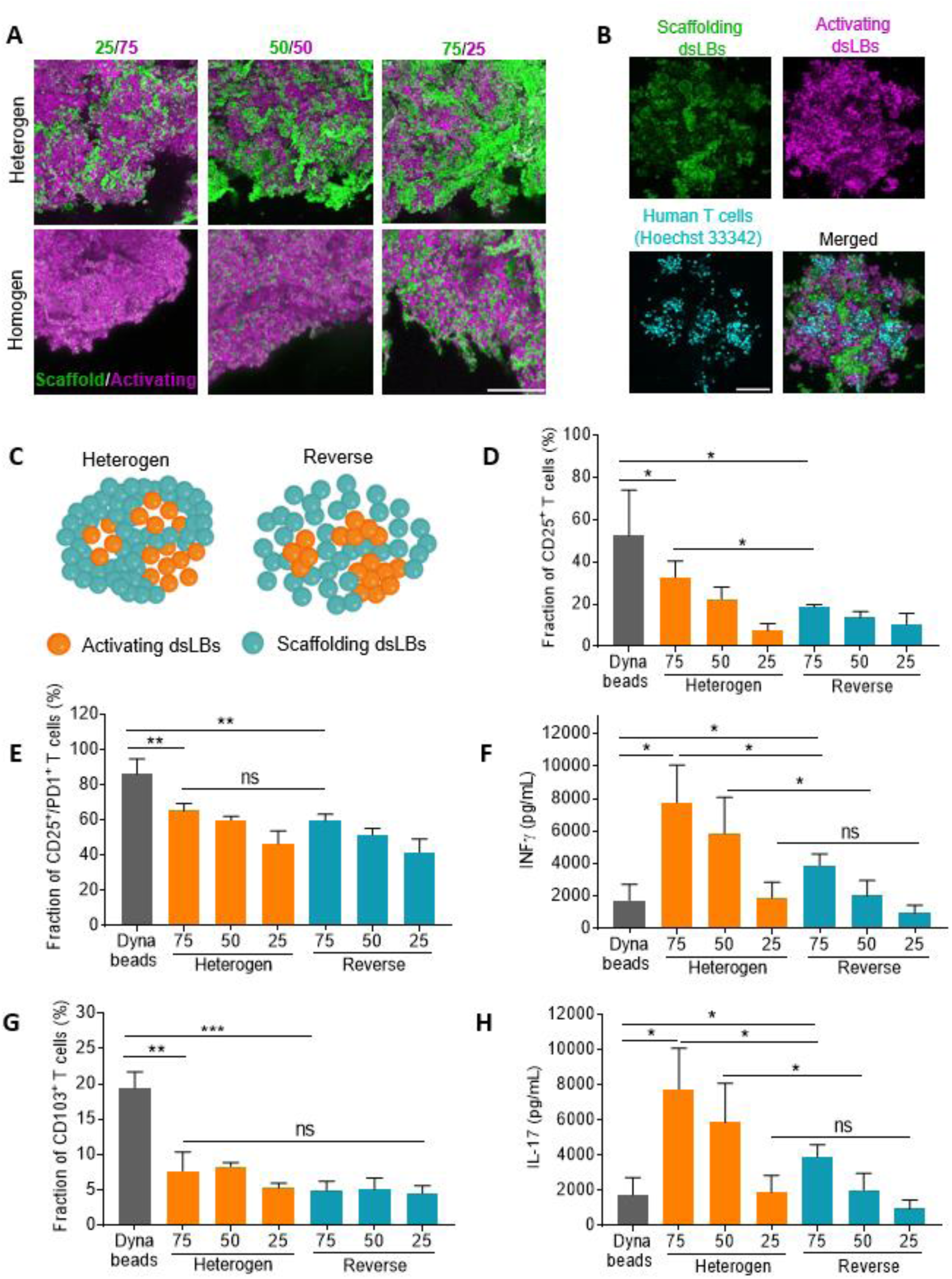
Heterogenous tissue formation and functionality. **A)** Representative confocal microscopy maximal z-projection of hetero-(top) and homogeneous (bottom) lymphBUTs formed from different ratios of activating dsLBs (magenta) and scaffolding dsLBs (green). Scale bar is 100 µm **B)** Representative confocal microscopy maximal z-projection of a heterogenous lymphBUT with immune-activation zones (magenta) and scaffold areas (green), incubated with human CD8^+^ T cells for 4 days and stained with Hoechst 33342 (cyan). The T cells expanded within the lymphBUT after 4 days of co-cultivation. Scale bar is 100 µm. **C)** Schematic illustration showing different lymphBUT constellations (heterogenous and reverse) assembled form 5 % and 1 % biotin dsLBs displaying no (blue) or a high anti-CD3 and anti-CD28 concentration (orange). **D) and E)** Flow cytometry quantification of CD25^+^ CD8^+^ T cell population (D) and CD25^+^/PD-1^+^ T cell subpopulation (E). **F)** Quantification of INFγ cytokine release *via* ELISA. **G) and H)** Phenotyping of human CD8^+^ T cells by quantifying the CD103 expression *via* flow cytometry (G) as well as analyzing the IL-17 (H) cytokine profile. Comparison of heterogenous and reverse lymphBUTs with varying ratios of activating and non-activating dsLBs (75 %, 50 %, 25 %) displaying a high anti-CD3 and anti-CD28 concentration with Dynabeads. Results are shown as mean +/- SD of two donors n > 2. P values were calculated using two-tailed t test. ns = not significant p>0.05, * p<0.05, ** p<0.01, *** p<0.001, **** p<0.0001

To assess directed migration of T cells into the activating zones of heterotypic lymphBUTs, we performed LSCM imaging of T cell infiltrated into the lymphBUTs after 4 days of co-cultivation and found T cell expansion clusters prevalently located in the activating areas rather than the scaffold (**Fig. 3 B**). Towards quantifying the relevance of lymphBUT heterogeneity and density for T cell expansion, we also engineered reverse lymphBUTs composed of 1 mol% T cell activating zones, surrounded by a denser 5 mol% non-functionalized scaffold (**Fig. 3 C**). We then measured expression markers and cytokine profiles of T cells incubated in the various lymphBUT configurations and compared these to Dynabeads.

Quantification of CD25^+^ T cells showed the highest T cell activation by Dynabeads of about 52.5 % followed by 75 heterogenous lymphBUTs with 32.6 % (**Fig. 3 D**). While reverse lymphBUTs have a lower ability to activate T cells compared to the heterotypic and homogenous conditions (**Fig S. 3 A**). This indicates the relevance of a looser scaffold serving as expansion matrix for T cell activation. For all conditions, a clear trend in CD25 expression depending on the concentration of activating dsLBs was observed (**Fig. 3 D, Fig S. 3 A**). A similar trend was detected when quantifying CD25^+^ PD-1^+^ T cells. PD-1 is an immune-suppression receptor that is usually upregulated by state-of-the-art *ex vivo* T cell expansion technologies. This upregulation diminishes the ability of cytokine secretion and is disadvantageous for a successful T cell therapy [20]. We confirmed a high fraction of CD25^+^ PD-1^+^ T cells activated with Dynabeads. Importantly, the PD-1 expression was significantly reduced for lymphBUT-expanded T cells (**Fig. 3 E, Fig S. 3 B**). Most importantly, the INFγ secretion, a critical effector cytokine that determines the efficacy of T cell therapies, revealed a significantly increased release by T cells expanded in lymphBUTs compared to Dynabeads (**Fig. 3 F, Fig S. 3 C**). This shows that while the overall activation efficacy (CD25) is lower in lymphBUTs, the expression of immuno-suppressive receptors (PD-1) and the secretion of effector cytokines (INFγ) is differentially regulated in the 3D expansion matrix pointing to a T cell phenotype with regulatory features beneficial for immunotherapy.

This observation was further supported by the quantification of CD103 expression, a tissue resident T cell marker, and secretion of the cytokines IL-10 and IL-17. The results demonstrate a significantly increased T cell population expressing CD103 when cultivated with Dynabeads compared to every other lymphBUT condition (**Fig. 3 G, Fig S. 3 D**). Contrary, the release of the pro-inflammatory cytokine IL-17 and the regulatory cytokine IL-10 is upregulated in T cells stimulated with lymphBUTs compared to Dynabead-expanded T cells (**Fig. 3 H, Fig S. 3 E, Fig S. 3 F**). Again, the lower the concentration of activating dsLBs incorporated into lymphBUTs the lower the cytokine concentrations detected. The results suggest that a considerable T cell activation can be achieved with the lymphBUT technology, offering a platform for expansion of a distinct CD8^+^ regulatory T cell phenotype as compared to the gold standard DynaBeads.

### Integrating metabolic active synthetic cells into lymphBUTs

Natural tissues not only regulate physiological functions *via* structural and mechanical properties but also provide an appropriate metabolic milieu. For instance, dendritic cells reduce the local pH in lymph nodes to sustain T cell differentiation and secrete H_2_O_2_ to further boost T cell expansion [21]. Towards integrating these metabolic aspects in lymphBUTs, we introduced metabolically active components in the form of enzyme-loaded unilamellar vesicles (UVs). LSCM confirmed the integration of a varying number of UVs, containing biotin-modified lipids in their membrane, into lymphBUTs (**Fig. 4 A and Fig S. 2 C**). To mimic the pH reduction and H_2_O_2_ production by dendritic cells, we loaded the UVs with glucose oxidase (GOx) enzymes and produced the UV with a membrane composition that allows for passive diffusion of glucose but retention of the enzyme [22]. GOx reduces the culture pH via glucose oxidation and produced H_2_O_2_ as a side product (**Fig. 4 B**).

**Fig. 4.**
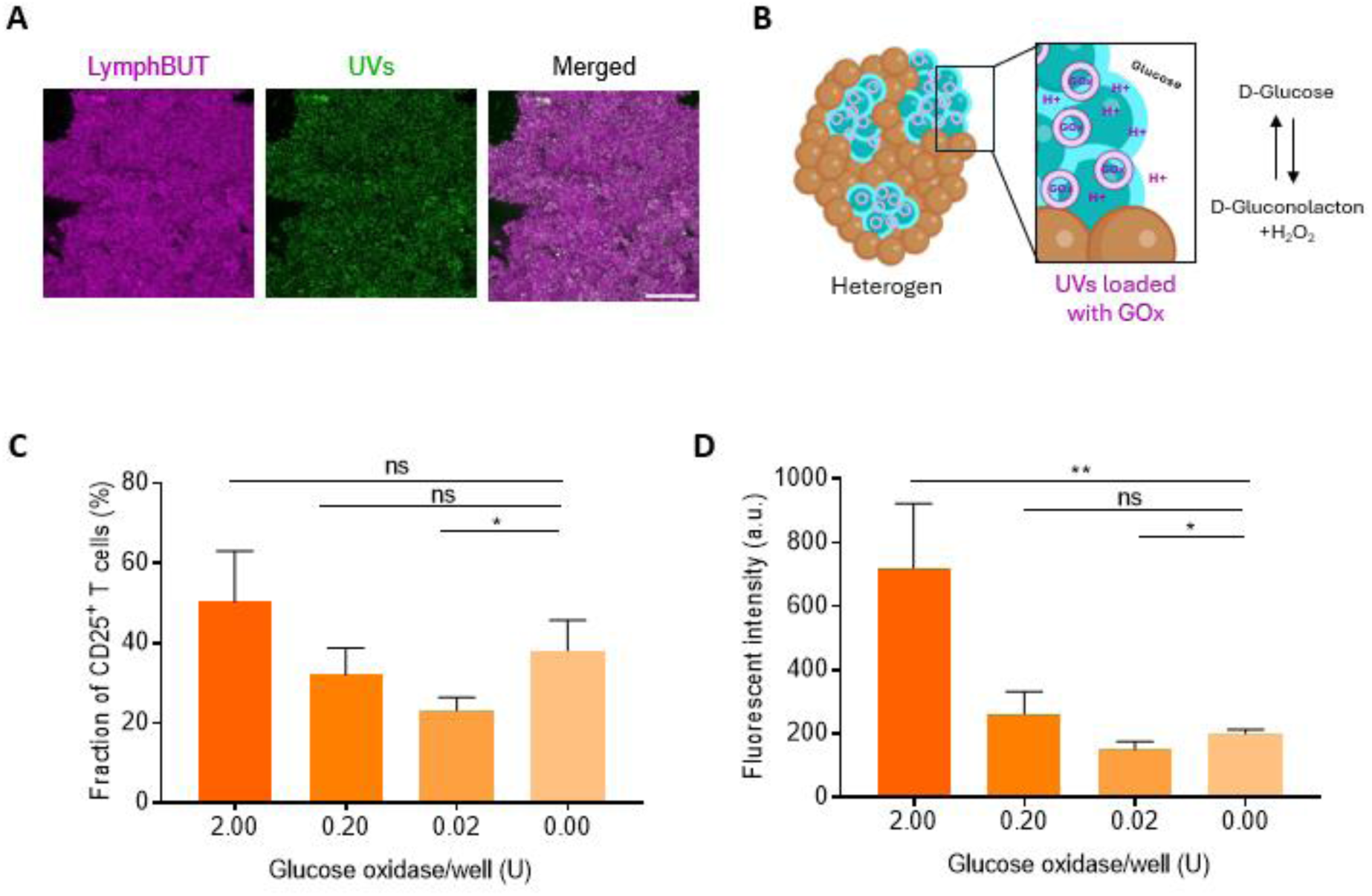
Integration of metabolic active components within lymphBUTs. **A)** Representative confocal microscopy image of UVs incorporated within a lymphBUT. Scale bar is 100 µm. **B)** Schematic illustration of glucose oxidase loaded UVs incorporated in the T cell activation zones of 75 % heterogenous lymphBUTs. **C) and D)** Flow cytometry quantification of CD25^+^ CD8^+^ T cell population (C) and metabolic T cell activity measured by Alamar blue assay (D) dependent on the UV concentration and therefore Glucose oxidase concentration in lymphBUTs (75 % heterotypic). Results are shown as mean +/- SD of two donors n = 2. P values were calculated using two-tailed t test. ns = not significant p>0.05, * p<0.05, ** p<0.01

By tuning the number of GOx-loaded UVs integrated into the 75/25 heterogeneous lymphBUTs, we produced synthetic tissues with 0.00, 0.02, 0.20 and 2.00 U GOx and cultivated these with primary human CD8^+^ T cells. The effect of GOx-modulated microenvironment within the synthetic tissues on T cell activation, as quantified by CD25 expression, and their metabolic activity, as assessed by resazurin reduction, was measured. While we did not resolve relevant differences in T cell activation, their metabolic activity was significantly increased in a GOx-dependent manner (**Fig. 4 C, D**). We detected an increase in metabolic activity of up to 3.5-fold for cultures with 2.00 U GOx. Interestingly, low concentrations of GOx significantly reduced T cell activation and metabolic activity which could be related to the increased activation zone density created by the UVs, limiting T cell infiltration and therefore activation. With increasing GOx, reduced pH and increased hydrogen peroxide concentrations, this effect could be overcome. The increase metabolic activity of T cells expanded with higher GOx again indicates a distinct phenotypic state of the cells, which is closely linked to their differentiation status [23]. Taken together, these results demonstrate that lymphBUTs are not only able to replicate tissue structure and mechanical dynamics but also integrate key biochemical triggers relevant to tune T cell expansion.

### Summary and Outlook

In this study we present a bottom-up synthetic biology inspired system for synthetic tissue assembly, meeting a biomedical need in immunotherapy. LymphBUTs are based on monodispersed synthetic cells, engineered to self-assemble and adding a third dimension to synthetic cell engineering [4]. With sizes of 1 – 4 mm they mimic structural and functional features of many natural tissues in relevant size ranges. Their overall density, architecture as well as their immunological and metabolic properties can be adjusted to mimic the structure and function of natural lymphatic tissues in a reduction of complexity approach.

We showcase their application for expansion of distinct T cell phenotypes in lymph node-mimicking environments. Previous technologies for this, are either based on monodispersed bead systems, focusing on receptor stimulation or artificial 3D matrixes integrating a chemical stimulus into bio-printed isotropic bulk materials [10]. In contrast, lymphBUTs can be formed with spatially segregated T cell activation zones, comprised of dense immune functionalized artificial antigen presenting cells, surrounded by an expandable scaffolding tissue and functionalized with metabolically active synthetic cells. With this, the synthetic tissues formed in lymphBUTs recreates three distinct features of tissues: Mechanical adaptability, metabolic activity and microstructural hierarchical organization.

We benchmarked the lymphBUT technology to commercial *ex vivo* T cell activation beads (Dynabeads). Flow cytometry and secretome results revealed the expression of a distinct regulatory CD8*^+^* T cell phenotype characterized by a high INFγ release as well as IL-10 and IL-17 secretion. This phenotype has already been reported and initial applicability for immunotherapeutic approaches has been suggested [16]. Even though the T cell activation measured by the CD25 signal is slightly lower on T cells cultivated with lymphBUTs as compared to DynaBeads, they showed a significantly reduced PD-1 expression, indicating a phenotype that is less prone to immune suppression. Interestingly, we observed differences in the activation dynamics comparing individual lmpyhBUT architectures. In natural lymphatic tissue the reaction areas, e.g. T cell zones or germinal center, host the majority of immune cells and feature dense networks of dendritic cells. In analogy to natural lymph nodes, lymphBUTs with denser activation zones and looser scaffolds were designed. By comparing T cell expansion in these lymphBUTs with reverse lymphBUTs we could show that this structure is crucial for efficient T cell expansion and demonstrate the bottom-up formation of a synthetic cell-based tissue with functional microanatomy.

Therefore, we successfully apply a synthetic biology-driven reduction of complexity approach to introduce structural cellular hierarchy in a synthetic tissue. In a future perspective, the co-cultivation of synthetic tissue with natural cells is not limited to immune cells and immunotherapeutic applications. It also provides a platform to mimic other tissue structures such as neurological tissue or hormone glands and study tissue signaling process in health and disease under defined molecular control in 3D.

### Experimental section

#### DsLB production

DsLBs were produced using electrostatic interactions in a layer-by-layer formation approach based on an oil in water emulsion. For that 100 mg of PDMS (Sylgard 184, Dow Corning USA) were pre-emulsified by resuspension in 910 µL of PBS adjusted to pH 7 with HCl and containing 4 mM of Sodium dodecyl sulfate (SDS, Sigma–Aldrich, Germany)). The oil in water droplets were emulsified for 2 min using a sonification bath. In a next step, 22 mM of MgCl_2_ (Sigma–Aldrich, Germany) and 100 µL of 6 mM small unilamellar vesicles (SUVs) were added and the mixture was incubated for 2 min protected from light. The excess SUVs were removed by centrifugation at 10000 xg for 30 sec, discharging of the supernatant and resuspension in 1 mL PBS adjusted to pH 7 with HCl. This washing step was repeated for a second time.

The SUVs were produced using extrusion as described previously [11]. They consist of 20 mol% EggPG, 5 mol% 18:1 MPB PE (maleimide), 1 mol% or 5 mol% 18:1 Biotinyl PE (biotin), 1 mol% LissRhodamine B-PE, Atto488-PE or 18:0 Cy5 PE and EggPC acting as a filling lipid (all Avanti Polar Lipids, USA).

#### DsLB immune functionalization

To integrate immune activating properties within the dsLB membrane a functionalized lipid equipped with a maleimide headgroup (Avanti Polar Lipids, USA) was integrated with 5 mol%. The maleimide headgroup can form a thiol connection with cysteine residues located on antibodies and proteins. For this antibody decoration, the total accessible amount of maleimide of the dsLB suspension needs to be determined. In order to do so, a SUV 1:1 dilution series ranging from 60 µM to 0.47 µM was prepared in a 96-well plate and used for calibration. The dsLB suspension was diluted 1:10 with PBS into the well plate.

The fluorescent intensity was measured with a TECAN Spark plate reader (Tecan Group, Switzerland) controlled by TECAN SparkControl software with in-built gain optimization. The excitation/emission settings were adjusted to Rhodamine B (537/582 nm), Atto488-PE (495/545 nm) or CY5-PE (630/675 nm) labeled dsLBs.

Based on the total lipid membrane concentration, the dsLBs were incubated with immune-stimulating antibodies against human CD3 (UCHT1, Invitrogen) and CD28 (CD28.2, Invitrogen) at a ratio (maleimide/antibody) of 0.5 for the high antibody condition and 0.125 for the low antibody condition and an adjusted αCD3/αCD28 ratio of 1:3. Before adding the antibody mix to the dsLBs, the antibodies were washed using the MircoSpin^TM^ G-25 Columns (Cytiva, US) according to manufacturer instructions. The antibody yield of 91 % (+/-1 %) was considered for the calculation (data not shown). The dsLBs were incubated with the antibodies in PBS adjusted to pH 7 at RT for 1 h resuspending every 15 min. The incubation was followed by a washing step. In order to know the dsLB concentration, the total lipid concentration was determined again by measuring the fluorescent intensity with the plate reader as already described.

#### MESF bead assay

To quantify the antibody concentration on the dsLB membrane a quantitative image-based Molecules of Equivalent Soluble Fluorochrome (MESF) assay was performed. For that dsLBs with 5 mol% maleimide in the membrane were produced and the lipid concentration was determined as described in dsLB immune functionalization. Based on the lipid concentration the accessible maleimide concentration was calculated. The dsLBs were functionalized with different ratios of maleimide/Alexa Fluor 488 anit-CD3 (HIT3a, BioLegend, UK) (1:1, 1:2, 1:4, 1:8, 1:16, 1:32). The Alexa Fluor 488 anit-CD3 was cleaned using the MircoSpin^TM^ G-25 Columns (Cytiva, US) according to manufacturer instructions and the antibody yield of 91 % (+/-1 %) was considered within the calculations. The mixture was incubated in PBS pH 7 for 1 h at RT protected from light and resuspended gently every 15 min. After the incubation the dsLBs were centrifuged at 10000 xg for 30 sec and resuspended in 200 µL PBS + 1 % BSA. The antibody decorated dsLBs were transferred to an 8-well Nunc LabTeK glass bottom chamber with mixture of Quantum Alexa 488 MESF beads (Bangs Laboratories Inc., USA) according to manufacturer instructions. The samples as well as the 488 MESF beads were imaged using a laser scanning microscope LSM 880 (Carl Zeiss AB) equipped with 20x objective and a 488 nm laser line. The analysis war performed with the ImageJ software (NIH, USA) by background subtraction, global-threshold segmentation, fill hole function, watershed particle separation and automated particle detection, measuring the fluorescent intensity of disperse dsLBs. The calibration was based on the four bead populations with known MESF/bead correlated to their fluorescent intensities. For calculating the antibody density per µm^2^ based on the MESF/area on dsLBs the fluorophores/antibody ratio was determined with the help of the NanoDrop (Thermo Fisher, Germany).

#### LymphBUT formation based on dsLBs

The incorporation of biotinylated lipids within the dsLB membrane combined with external provided streptavidin allows the self-assembly of lymphBUTs. For this purpose, dsLBs were produced as already described with additional 1 mol% or 5 mol% Biotin (Avanti Polar Lipids, USA) in the lipid membrane. In order to form lymphBUTs of about a similar size and number of dsLBs, the total lipid concentration of the dsLB membrane was determined as described above by measuring the fluorescent intensity using the TECAN Spark plate reader (Tecan Group, Switzerland). A total number of approximately 2.7 million dsLBs were transferred to a 48-well plate. These wells were filled with PBS to a final volume of 400 - 450 µL containing streptavidin (Invitrogen, USA) in a molecular biotin/streptavidin ratio of 50, 100, 200, 400, 800, 1600, 3200 or 6400:1. By orbital shaking at 450 rpm for 1-2 h the biotin was exposed to the diffusing streptavidin leading to the lymphBUT self-assembly.

#### LymphBUT architectures

The lymphBUT formation strategy based on a strong connection between biotin and streptavidin allows the engineering of diverse lymphBUT architectures. LymphBUTs can be build form dsLBs displaying no T cell activating antibodies (plain). Contrary, they can be formed of dsLBs decorated with immune-stimulating antibodies against human CD3 (UCHT1, Invitrogen) and CD28 (CD28.2, Invitrogen) in a 1 to 3 ratio and high (420 molecules/µm^2^) or low antibody conditions (320 molecules/µm^2^).

In order to achieve a hierarchical structure within a heterotypic lymphBUT a sequential formation procedure was developed. In a first step, a lymphBUT was formed form T cell activating, high antibody decorated 5 % biotin dsLBs using a biotin/streptavidin ratio of 400:1 in 48 well plates with a final PBS volume of 400 µL. After 1 h of orbital shaking at 450 rpm the lymphBUT was well formed and 200 µL of PBS were disposed. The formed lymphBUT was resuspended gently in the remaining PBS, thus leading to smaller lymphBUTs of sizes between 50 – 100 µm later acting as T cell activation zones. In a second step, 1 % biotin non-functionalized dsLBs were added with the appropriate streptavidin concentration of 400:1 biotin/streptavidin ratio to form a connecting scaffold around the already existing, smaller T cell activating lymphBUT zones.

Vise versa to the heterotypic lymphBUTs, reverse heterotypic lymphBUTs were formed to investigate the impact on the activation zone and scaffold density on the activation dynamics and phenotyping of human CD8^+^ T cells. For this, the activation zones were built from 1 % biotin dsLBs with a high antibody functionalization surrounded by 5 % biotin non-functionalized scaffolding dsLBs.

In contrast to the heterotypic lymphBUT architectures, we formed homogenous lymphBUTs by pre-mixing 5 % biotin functionalized and 1 % biotin non-functionalized dsLBs connection with streptavidin of a total biotin/streptavidin ratio of 400:1.

The ratios of immune stimulating dsLBs to connective scaffold dsLBs were varied (75/25, 50/50 and 25/75). The total dsLB number was kept at approximately 2.7 million dsLBs per lymphBUT.

#### LymphBUT formation based on silica beads

To compare the lymphBUT formation from dsLBs with stiffer beads, silica microspheres (Bangs Laboratories Inc., USA) with a diameter of 3.94 µm were covered with a lipid bilayer membrane identical to the dsLB membrane. For that 25 µL of silica beads were resuspended in PBS and centrifuged at 10000 xg for 30 sec. The washing step war repeated a second time, and the beads were resuspended in 100 µL PBS before incubating for 2 min with 50 µL SUVs containing 20 mol% EggPG, 5 mol% 18:1 MPB PE, 5 mol% 18:1 Biotinyl PE, 1 mol% LissRhodamine B-PE and 69 mol% EggPC (all Avanti Polar Lipids, USA). After the SUV incubation the silica beads were washed twice and resuspended in 500 µL PBS. The wells of a 48-well plate were prepared with 400 – 450 µL of PBS supplemented with streptavidin (1:100, 1:400 and 1:1600 streptavidin/biotin ratio). The beads shook overnight at 400 µL in an orbital motion.

#### Stereomicroscopy imaging of lymphBUTs

For analyzing the self-assembly process of lymphBUTs with 1 mol% or 5 mol% biotin (Avanti Polar Lipids, USA) in the dsLB membrane and different streptavidin (Invitrogen, USA) concentrations in the PBS environment, lymphBUTs were produced as just described. They were visualized in the 48 well plate by bright field images acquired with a LEICA DFC450 stereo microscope (Leica Microsystems, Germany) and a 2x (PLANAPO 2.0x/39, Leica Microsystems, Germany) or a 5x objective (PLANAPO 5.0x/0.5, Leica Microsystems, Germany). The images were analyzed with the ImageJ software (NIH, USA) by background subtraction of the LUT inverted image, a global-threshold segmentation, fill hole function, watershed particle separation and automated particle detection. The parameter considered for the formation robustness of lymphBUTs was the mean intensity of the single lymphBUTs which was related to the density and therefore to the stability of the lymphBUTs and the robustness of the formation process.

#### Laser scanning confocal microscopy imaging of lymphBUTs

The lymphBUTs comprising activation zones connected with a non-functionalized dsLB scaffold was visualized using the confocal laser scanning microscope LSM 880 (Carl Zeiss AB) equipped with a 10x objective (EC “Plan-Neofluar” 10x/0.30 M27, Carl Zeiss AG, Germany) and 405, 488 and 647 laser lines. To visualize the human CD8^+^ T cell which infiltrated the lymphBUT, the T cell nuclei were fixed with 2 % PFA and stained with 1.5 nM Hoechst 33342 trihydrochloride (Thermo Fisher, Germany). The images were analyzed with the ImageJ software (NIH, USA) by z-projection and background subtraction.

#### Connection area analysis using LSCM

In order to compare the connection areas between single dsLBs, they were either equipped with 1 % or 5 % biotin and 1% Cy5 in the dsLB membrane. Those dsLBs were then connected to lymphBUTs with Alexa Fluor 405 labeled streptavidin (Thermo Fisher, Germany) at a ratio of 1400:1 biotin/streptavidin.

The resulting constructs were then carefully transferred to an 8-well Nunc LabTeK glass bottom chamber slide filled with 200 µL PBS. The lymphBUTs were imaged with the laser scanning microscope LSM 880 (Carl Zeiss AB) equipped with a 63x immersion oil objective (Plan-Apochromat 63x/1.4 oil DIC M27, Carl Zeiss AG, Germany) and 405 and 647 nm laser lines. The images were analyzed with the ImageJ software (NIH, USA) by background subtraction and manual selection of the connection areas.

#### Mechanical properties of lymphBUTs

The mechanical properties of lymphBUTs were determined by micro indentation. For that dsLBs with 1 % and 5 % biotin in the membrane were connected with 400:1 biotin/streptavidin ratio and measured with the Microindenter G2 CellScale (CellScale biomaterials testing, Canada). To do so, the lymphBUTs were transferred to the CellScale fluid bath filled with 45 mL PBS on top of the testing anvil. The tungsten microbeam with a diameter of 0.0762 mm and a modulus of 411000 MPa was equipped with a 1×1 mm square stainless-steel platen functioning as cantilever. The mechanical properties of lymphBUTs were measured by z compression with a compression magnitude of 10 % of the lymphBUT height. The microbeam displacement was controlled in a ramp setting with a loading duration of 30 sec, a holding duration of 5 sec and a recovery duration of 30 sec. The analysis is based on real time image tracking at a frequency of 5 Hz. The images were acquired using a USB digital camera with a zoom lens and XYZ automatized stage. The indentation force for lymphBUTs produced in two individual batches with 1 % and 5 % biotin in the dsLB membrane was measured (n=9).

#### Integrating metabolic components into the lymphBUT

As many natural tissues show metabolic features, metabolic components were integrated into lymphBUTs in the form of enzyme loaded unilamellar vesicles (UVs). To produce the LUVs 5 mol% 18:1 Biotinyl PE (biotin), 1 mol% LissRhodamine B-PE, Atto488-PE and 94 mol% EggPC (all Avanti Polar Lipids, USA). The lipids were pre-mixed and dried for 1 h using a vacuum pump. A sucrose (Sigma Aldrich, UK) solution diluted to 285 mM with PBS and 2000 U Glucose Oxidase from Aspergillus niger (Sigma Aldrich, UK) per mL were added to the dried lipids and incubated for 15 min protected from light. After the incubation the lipids were resuspended and extruded using Nucleopore^TM^ membrane of 1.0 µm pores (Cytiva, Germany). In order to get rid of the excess Glucose Oxidase, the UVs were diluted 1:200 with PBS and centrifuged at 10,000 xg for 6 min. The supernatant was deposed, and the pellet was resuspended in fresh PBS. The washing step was repeated three times. The LUVs were integrated into the 5 % biotin lymphBUT activation zones resulting in a total Glucose oxidation concentration in the lymphBUTs of 0.02, 0.20 and 2.00 U.

#### T cell isolation and cultivation

Primary human CD8^+^ T cells have been used for functional assays with lymphBUTs. The T cells were isolated from leukapheresis reduction system chambers from healthy voluntary blood donors. Negative isolation was performed with commercial selection kits (RosetteSep Human CD8^+^ T cell Enrichment Cocktail, STEMCELL technologies, Germany) following the manufacturers instruction. The Institute for Clinical Hemostaseology and Transfusion Medicine, Saarland University Medical Center provided the blood following ethics agreement number 34/23 (Ethikkommission Ärztekammer des Saarlandes). After the isolation the T cells were frozen in RPMI 1640 w/ L-Glutamine (VWR, Germany) medium supplemented with 40 % fetal bovine serum (Gibco, Germany), 1 % penicillin/streptomycin (Gibco, Germany), 1 % non-essential amino acids (Biowest) and 50 mM HEPES (Sigma–Aldrich, Germany) and additional 10 % dimethyl sulfoxide (DMSO) (Sigma Aldrich, Germany) and stored at -80 °C until further use.

The primary human CD8^+^ were thawed one day before starting the experiment and incubated overnight. The T cells were cultivated in RPMI 1640 w/L-Glutamine (VWR, Germany) medium supplemented with 10 % fetal bovine serum (Gibco, Germany), 1 % penicillin/streptomycin (Gibco, Germany), 1 % non-essential amino acids (Biowest, France) and 50 mM HEPES (Sigma–Aldrich, Germany) with 50 U mL^-1^ of the growth factor Interleukin 2 (IL-2) in T 25 cell culture flasks at 37 C and 5 % CO_2_.

#### T cell *ex vivo* activation and expansion within lymphBUTs

The isolated primary human CD8^+^ T cells were co-cultivated with lymphBUTs in round bottom 96 well plates and fully supplemented RPMI 1640 medium. Therefore, different lymphBUT architecture as described were formed, washed and transferred with 100 µL to the round bottom 96 well plates. Dispersed high antibody functionalized dsLBs and Dynabeads human T-Activator CD3/CD28 beads (Gibco, Germany) used according to manufacturer suggestions served as controls.

Because the lymphBUTs cannot form in RPMI medium due to the supplementation of vitamins including biotin they were formed in PBS. Therefore, they need to be extensively washed with fully supplemented RPMI 1640 medium before transferring within 100 µL of medium to the round bottom 96 well plate. Because of the lymphBUT size of 1-4 mm, 1/10 of the pipette tip need to be cut to increase the diameter of the tip opening and not destroy the lymphBUT. 80.000 T cells were transferred to each well containing a lymphBUT leading to a total medium volume of 250 µL and IL-2 concentration of 50 U mL^-1^. The wells at the outer border were filled with PBS in order to avoid evaporation and therefore biased results. The primary human CD8^+^ T cells were incubated with the lymphBUTs in fully supplemented RPMI 1640 medium at 37 °C and 5 % CO_2_ for 4 days.

#### Cytokine analysis using an automated ELISA system

In order to analyze T cells activation, expansion as well as T cell phenotyping, primary human CD8^+^ T cells were co-cultivated with different lymphBUTs architectures in fully supplemented RPMI1640 medium for 4 days. After the incubation the 80 µL of supernatant were transferred to a another 96 well plate and either used directly or frozen at -80 °C until further use. The cytokine concentrations of IL-10, IL-17A and IFNγ 3^rd^ gen were measured with the Ella^TM^ Automated ELISA protein simple system (Biotechne, USA). The samples were prepared according to manufacturer instructions and analyzed with the Simple Plex Runner software (Biotechne, USA).

#### CD8^+^ T cell activation studies using Flow Cytometry

To further study primary human CD8^+^ T cell activation and immunosuppression signals as well as phenotype related surface receptors, the remaining lymphBUTs hosting the CD8^+^ T cells were resuspended, centrifuged at 300 xg for 5 min, the supernatant was disposed and the cells were resuspended in PBS containing staining antibodies (1:400) against the surface marker CD25 (BC96, BioLegend, UK), PD-1 (NAT105, BioLegend, UK) and CD103 (Ber-ACT8, BioLegend, UK) conjugated to Alexa Fluor 488 or Alexa Fluor 647. The cells were incubated with the antibody solution for 1 h at RT protected from light. After incubation the cells were washed once and resuspended in PBS containing 2 % PFA (Sigma–Aldrich, Germany) for 30 min at RT protected from light in order to fix the CD8^+^ T cells. The cells were than washed again to remove the remaining PFA and resuspended in PBS containing 1.5 nM Hoechst 33342 trihydrochloride (Thermo Fisher, Germany) for staining the T cell nucleus for 20 min protected from light. Finally, the cells were washed once more and resuspended in PBS + 1 % Albumin fraction V (BSA) (Sigma–Aldrich, Germany). For the quantification of the surface marker the Attune NxT Flow Cytometer and the Attune^TM^ Software (Thermo Fisher, Germany) were used. The flow cytometer is equipped with 405, 488, 561 and 637 nm laser lines. For the analysis a minimum of 10,000 events were considered and analyzed with the FlowJo V.10 software (FlowJo LLC, USA).

#### Validating the human CD8^+^ T cell metabolic activity *via* Alamar blue assay

In order to compare the metabolic activity of human CD8^+^ T cells co-cultivated with metabolic lymphBUTs containing UVs loaded with varying glucose oxidase concentrations an alamar blue assay was performed. For that reason, 10 v% of the alamarBlue solution (Bio Rad, UK) was added after 4 days of co-cultivation. The cells were then incubated for 4 h at 37 °C. The living T cell will convert resazurin to resorufin. The fluorescent resorufin is than measured at the TECAN Spark plate reader (Tecan Group, Switzerland). The excitation/emission settings were adjusted to 560/560 nm with a bandwidth of 10 nm. The measured fluorescent intensity relative to metabolic activity and therefore indirectly to the T cell proliferation.

### Data processing and statistical analysis

Graphs were plotted with GraphPad Prism 8 as mean +/- SD of technical and biological replicates. Statistical analyses were performed with the in-build GraphPad Prism 8 software function. The applied statistical analysis were noted for the individual figures in the figure legends. Schematic illustrations were created with BioRender.com.

## Acknowledgements

The authors thank Nils Piernitzki, Alpcan Önür, the INM Fluorescence Microscopy Core Facility and Cao Nguyen Duong (Leibniz Institute for New Materials) for their expertise in dsLB production, micro indentation measurements and confocal microscope use. We also thank Kathleen Seelert (CIPMM) for her help with blood and T cell handling and collection. The authors acknowledge funding from the Pharmazeutische Forschungsallianz Saarland, the Daimler and Benz Foundation (32-12/22), the Joachim Herz Foundation (Add-on Fellowship and Innovate! Akademie) and German Science Foundation (Emmy Noether Program, project number 525255627 and 545610076).

## References

[1] H. Bayley, I. Cazimoglu, C.E.G. Hoskin, Synthetic tissues, Emerg Top Life Sci 3(5) (2019) 615–622.

[2] F. Lussier, O. Staufer, I. Platzman, J.P. Spatz, Can Bottom-Up Synthetic Biology Generate Advanced Drug-Delivery Systems?, Trends Biotechnol 39(5) (2021) 445–459.

[3] X. Yan, X. Liu, C. Zhao, G.Q. Chen, Applications of synthetic biology in medical and pharmaceutical fields, Signal Transduct Target Ther 8(1) (2023) 199.

[4] Ö.D. Toparlak, Artificial cells drive neural differentiation, Science Advances 6(38) (2020).

[5] J.E. Hernandez Bucher, O. Staufer, L. Ostertag, U. Mersdorf, I. Platzman, J.P. Spatz, Bottom-up assembly of target-specific cytotoxic synthetic cells, Biomaterials 285 (2022) 121522.

[6] N. Hakami, A. Burgstaller, N. Gao, A. Rutz, S. Mann, O. Staufer, Functional Integration of Synthetic Cells into 3D Microfluidic Devices for Artificial Organ-On-Chip Technologies, Adv Healthc Mater 13(22) (2024) e2303334.

[7] A. Dupin, F.C. Simmel, Signalling and differentiation in emulsion-based multi-compartmentalized in vitro gene circuits, Nat Chem 11(1) (2019) 32–39.

[8] W.K. Spoelstra, S. Deshpande, C. Dekker, Tailoring the appearance: what will synthetic cells look like?, Curr Opin Biotechnol 51 (2018) 47–56.

[9] G. Villar, A.D. Graham, H. Bayley, A tissue-like printed material, Science 340(6128) (2013) 48–52.

[10] A. Alcinesio, I. Cazimoglu, G.R. Kimmerly, V. Restrepo Schild, R. Krishna Kumar, H. Bayley, Modular Synthetic Tissues from 3D-Printed Building Blocks, Advanced Functional Materials 32(7) (2021).

[11] A. Burgstaller, N. Piernitzki, N. Kuchler, M. Koch, T. Kister, H. Eichler, T. Kraus, E.C. Schwarz, M.L. Dustin, F. Lautenschlager, O. Staufer, Soft Synthetic Cells with Mobile Membrane Ligands for Ex Vivo Expansion of Therapy-Relevant T Cell Phenotypes, Small (2024) e2401844.

[12] K.H. Biswas, Molecular Mobility-Mediated Regulation of E-Cadherin Adhesion, Trends Biochem Sci 45(2) (2020) 163–173.

[13] D. Zhao, D. Zhu, F. Cai, M. Jiang, X. Liu, T. Li, Z. Zheng, Current Situation and Prospect of Adoptive Cellular Immunotherapy for Malignancies, Technol Cancer Res Treat 22 (2023) 15330338231204198.

[14] A.J. Najibi, D.J. Mooney, Cell and tissue engineering in lymph nodes for cancer immunotherapy, Adv Drug Deliv Rev 161–162 (2020) 42-62.

[15] H.S. Kim, T.C. Ho, M.J. Willner, M.W. Becker, H.W. Kim, K.W. Leong, Dendritic cell-mimicking scaffolds for ex vivo T cell expansion, Bioact Mater 21 (2023) 241–252.

[16] Y. Shou, S.C. Johnson, Y.J. Quek, X. Li, A. Tay, Integrative lymph node-mimicking models created with biomaterials and computational tools to study the immune system, Mater Today Bio 14 (2022) 100269.

[17] S.C. Johnson, J. Frattolin, L.T. Edgar, M. Jafarnejad, J.E. Moore, Jr., Lymph node swelling combined with temporary effector T cell retention aids T cell response in a model of adaptive immunity, J R Soc Interface 18(185) (2021) 20210464.

[18] M.L. Manning, R.A. Foty, M.S. Steinberg, E.M. Schoetz, Coaction of intercellular adhesion and cortical tension specifies tissue surface tension, Proc Natl Acad Sci U S A 107(28) (2010) 12517–22.

[19] N. Prokhnevska, M.A. Cardenas, R.M. Valanparambil, E. Sobierajska, B.G. Barwick, C. Jansen, A. Reyes Moon, P. Gregorova, L. delBalzo, R. Greenwald, M.A. Bilen, M. Alemozaffar, S. Joshi, C. Cimmino, C. Larsen, V. Master, M. Sanda, H. Kissick, CD8(+) T cell activation in cancer comprises an initial activation phase in lymph nodes followed by effector differentiation within the tumor, Immunity 56(1) (2023) 107–124 e5.

[20] J.W. Hickey, A.K. Kosmides, J.P. Schneck, Engineering Platforms for T Cell Modulation, Int Rev Cell Mol Biol 341 (2018) 277–362.

[21] M. Los, W. Droge, K. Stricker, P.A. Baeuerle, K. Schulze-Osthoff, Hydrogen peroxide as a potent activator of T lymphocyte functions, Eur J Immunol 25(1) (1995) 159–65.

[22] S. Purushothaman, J. Cama, U.F. Keyser, Dependence of norfloxacin diffusion across bilayers on lipid composition, Soft Matter 12(7) (2016) 2135–44.

[23] S. Ma, Y. Ming, J. Wu, G. Cui, Cellular metabolism regulates the differentiation and function of T-cell subsets, Cell Mol Immunol 21(5) (2024) 419–435.

