## Supplementary material for "Synthetic cell-based tissues for bottom-up assembly of artificial lymphatic organs": Fig S.

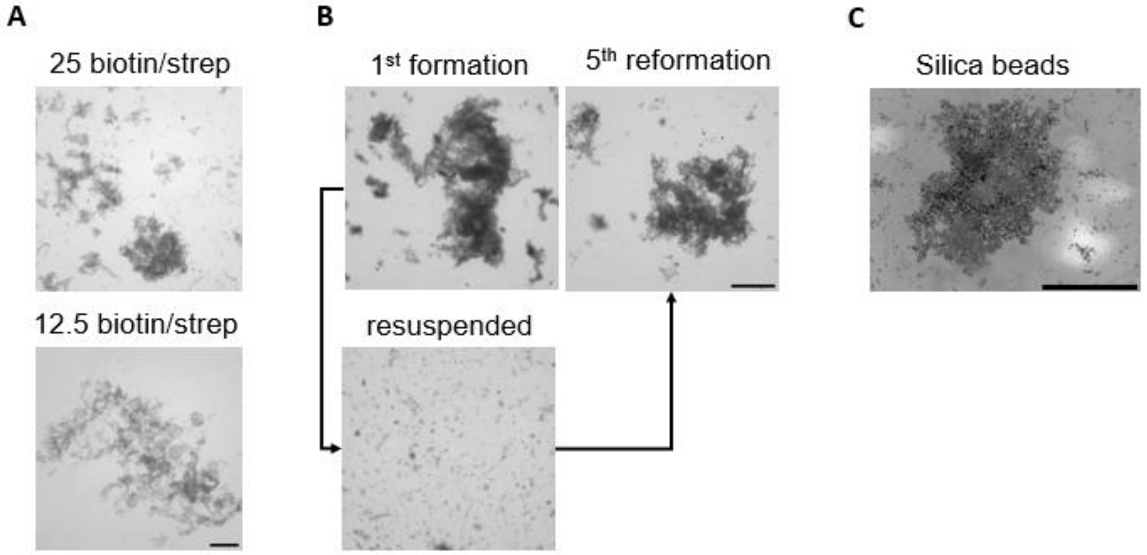


**Fig S. 1: LymphBUT formation A)** Stereo microscopy image of lymphBUTs formed with biotin/streptavidin ratios of 25 and 12.5:1. **B)** Stereo microscopy images showing the re-formation property of lymphBUT after resuspension over 6 cycles. **C)** Silica beads surrounded by a lipid bilayer as used for dsLBs interconnected *via* biotin/streptavidin bond at a ratio of 100:1. Scale bares are 1 mm.

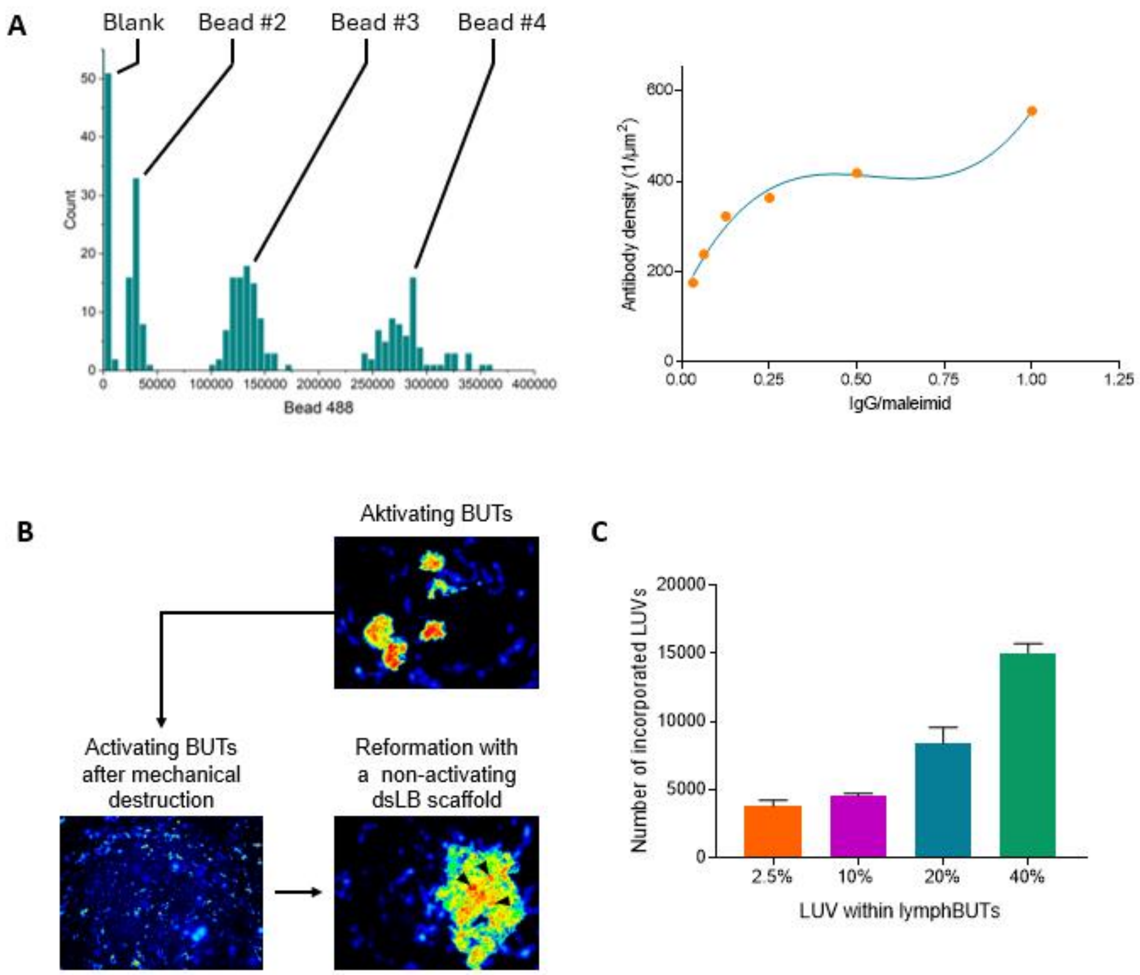


**Fig S. 2: Functional and structural lymphBUT properties A)** Fluorescent intensity quantification using MESF calibration beads for quantifying IgG density on dsLBs. **B)** Sequential self-assembly process for heterotypic lymphBUT formation. **C)** Number of incorporated LUVs in relation to the initially added concentration analyzed via confocal imaging.

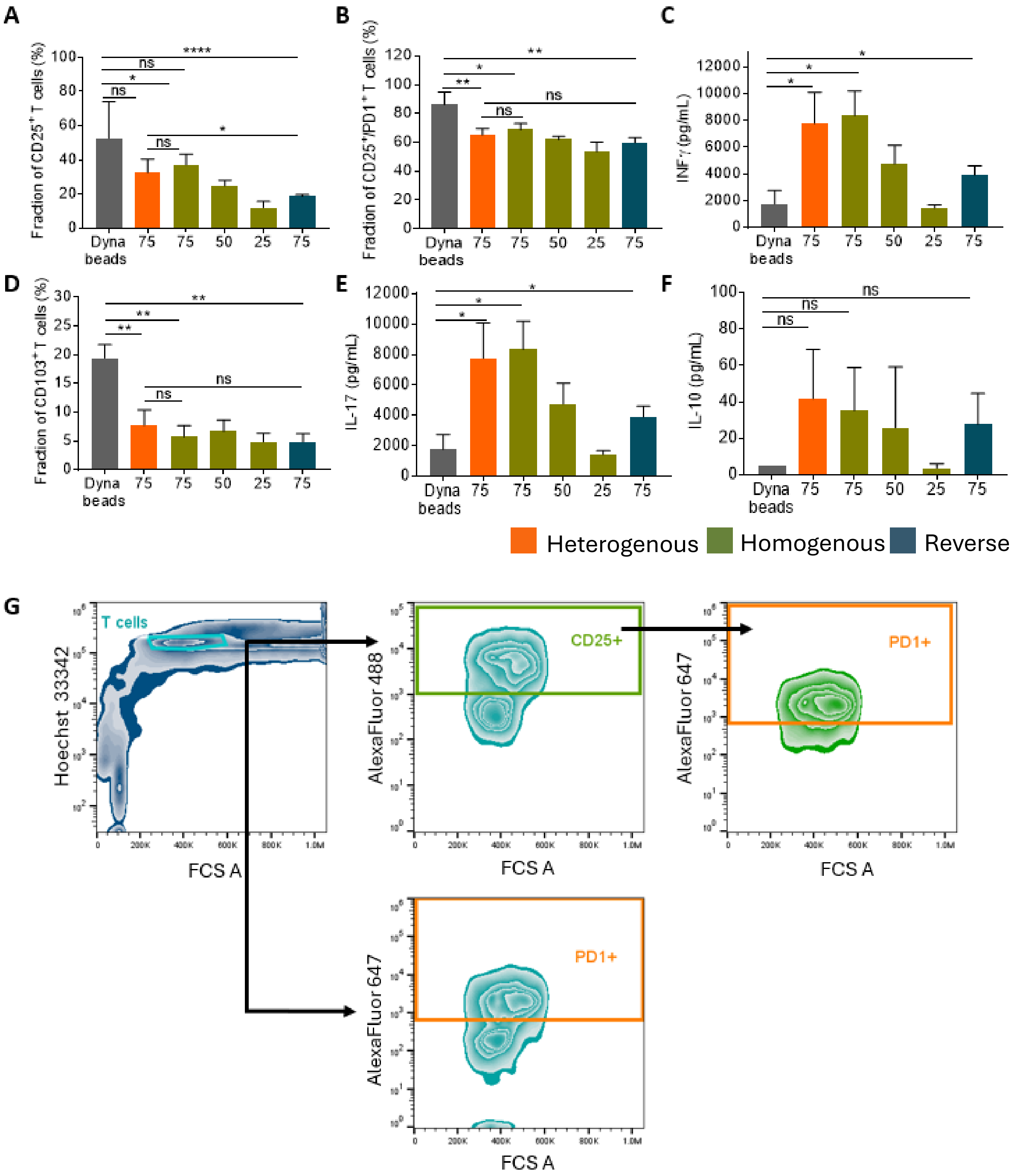


**Fig. S 3 Activation and immunosuppression profile of expanded T cells depending on the lymphBUT architectures A) and B)** Flow cytometry quantification of $CD25^+$ $CD8^+$ T cell population (B) and $CD25^+$/$PD\text{-}1^+$ T cell subpopulation (D). **C)** Quantification of INFγ cytokine release *via* ELISA. **D), E) and F)** Phenotyping of human $CD8^+$ T cells by quantifying the CD103 expression *via* flow cytometry (F) as well as analyzing the IL-17 (F) and IL-10 (H) cytokine profile. Comparison of heterogenous (orange), homogenous (green) and reverse (blue) lymphBUTs with varying ratios of activating and non-activating dsLBs (75 %, 50 %, 25 %) displaying a high anti-CD3 and anti-CD28 concentration with Dynabeads. **G)** Flow cytometry gating strategy shown using the example of the Dynabead control. Results are shown as mean +/- SD of two donors n > 2. P values were calculated using two-tailed t test. ns = not significant p>0.05, * p<0.05, ** p<0.01, **** p<0.0001
